# Practical considerations regarding estimating *N*_*e*_ when generations overlap

**DOI:** 10.1101/2024.06.04.597384

**Authors:** Robin S. Waples

**Author notes:** Corresponding author: Robin Waples.

## Abstract

Researchers studying species in nature often find it challenging to apply methods based on simplistic models of reality. Here I consider how some real-world complications influence demographic estimates of effective population size (*N*_*e*_) when generations overlap. The most widely-used model (by Hill) expresses *N*_*e*_ as a function of variance in lifetime reproductive output (*LRO*) of the *N*_1_ members of a newborn cohort. Hill’s model assumes stable age structure and constant population size, in which case mean 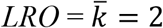. In real-world applications, researchers often ask whether unbiased estimates can be obtained under the following conditions: (1) When 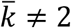 for empirical data; (2) If cohorts are defined at a later age than newborns; (3) If survival to age at sexual maturity (α) is not random; (4) When some or all null parents (those with *LRO*=0) are not sampled. Using analytical methods and computer simulations, I show that: (1) Because variance in offspring number is positively correlated with the mean, 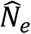 will be biased using raw data when 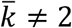, but this bias can be overcome by rescaling var(*LRO*) to its expected value when 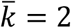. (2) The cohort can be defined at any age ≤α, provided that (a) *LRO* data cover the full lifespan (e.g., production of newborns by newborns, or production of adults by adults), and (b) survival to age α is random. (3) If juvenile survival is family-correlated, defining cohorts at age α avoids upward bias in 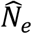 that occurs if newborn cohorts are used. (4) Missing some or all null parents has no effect on 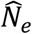, provided that data are rescaled to 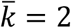.

## INTRODUCTION

The most widely-used demographic method for calculating *N*_*e*_ when generations overlap is by Hill (1972, 1979). If age-structure is stable and the population produces a constant number (*N*_*1*_) of offspring in each cohort, *N*_*e*_ per generation is

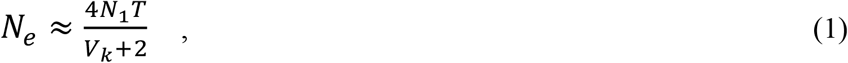

where *T* is generation length (average age of parents of a newborn cohort) and *V*_*k*_ is the variance in lifetime reproductive output (*LRO*) = variance in lifetime number of offspring among the *N*_*1*_ members of a cohort. Equation 1 was derived in terms of the variance in allele frequency and hence applies to the variance effective size. However, because the derivation assumes constant *N* (in which case mean 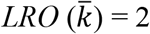), Equation 1 also gives the inbreeding effective size as well (Johnson 1977). In spite of the constant-*N* and stable-age-structure assumptions, Equation 1 is robust to random demographic variation in birth and death rates (Waples et al. 2014), and it accurately predicts *N*_*e*_ for extreme reproduction scenarios (strong reproductive skew and strong temporal correlations between individual reproduction and survival; Waples 2023).

Nevertheless, in real-world applications, researchers are often confronted with empirical data that appear to conflict with assumptions of Hill’s model. Here I consider four common scenarios and how they affect performance of 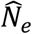 using Equation 1. (1) Even if population size is constant, it generally will not be the case that 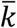 in a sample is exactly 2. Because the variance in offspring number is positively correlated with the mean (Crow and Morton 1955), this has the potential to bias 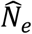 for estimates based on raw data. (2) Hill defined *N*_*1*_ to be the number of newborn individuals in a cohort. In many species, however, it is difficult or impossible to sample newborns and keep track of their lifetime production of offspring. How robust is Equation 1 to using a different age to define the cohorts? (3) What happens if survival of newborns to age at sexual maturity (α) is not random, and how does non-random survival interact with the age for defining the cohorts? (4) What happens if some or all null parents (those with *LRO* =0) are not available to be sampled? Here, these questions are addressed using a combination of analytical and numerical methods.

## METHODS

### Notation

The underlying demographic model used here assumes strongly-seasonal, birth-pulse reproduction as defined by Caswell (2001). Upon reaching age *x*, each individual produces on average *b*_*x*_ offspring and then survives to age *x*+1 with probability *s*_*x*_. Cumulative survival through age *x* is *l*_*x*_. Because age 0 individuals generally do not reproduce, offspring that survive to age 1 are considered ‘newborns’, and *b*_*x*_ is scaled to production of yearling offspring, and *l*_*x*_ is defined to be 1. Age at sexual maturity (α; α≥1) is assumed fixed. Sexes are separate and vital rates can differ between males and females.

Let *k*_*i*_ be *LRO* for the *i*^th^ member of a newborn cohort. Then 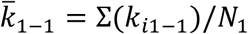, where the ‘1-1’ in the subscript indicates production of newborns by newborns (hence covering a full life cycle). From the definition of a variance,

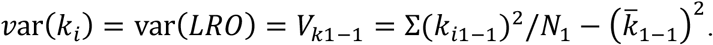

In theory, with constant 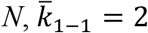 and *V*_*k*1−1_ = Σ(*k*_*i*1−1_)^2^/*N*_1_ − 4. More generally, we have to allow for the possibility that 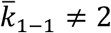

### Mean offspring number ≠ 2

Using a discrete-generation model, Crow and Morton (1955) considered a scenario where offspring of a highly-fecund species are sampled at an early life stage, where it is expected that 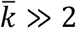. Except in the special case of a Poisson variance in offspring number, the variance is positively correlated with the mean, so *V*_*k*_ will generally be inflated when 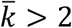. The solution Crow and Morton proposed was to rescale raw *V*_*k*_ to the value it would be expected to take if random mortality of offspring occurred until mean offspring number reached some smaller specified value. Mathematically, this is exactly equivalent to randomly subsampling the offspring.

Let the subscripts ‘raw’ refer to the initial empirical data and ‘Adj’ refer to adjusted values. Then Crow and Morton’s general rescaling method is (from Equation 3 in Waples 2002):

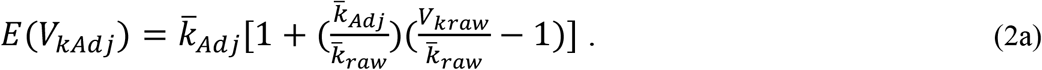

To rescale *V*_*kraw*_ to its expected value in a stable population, set 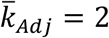:

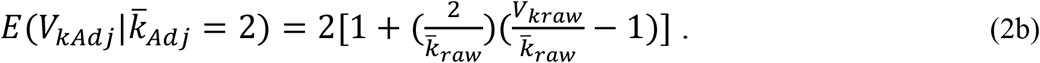

Although Crow and Morton (1955) only considered scenarios where 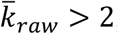, Equations 2a,b also correctly predict *V*_*kAdj*_ when 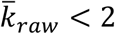 (Waples 2002), in which case *V*_*kAdj*_ is biased downwards compared to its stable-*N* expectation. In this latter case, *E*(*V*_*kAdj*_) is the variance one would expect to find if the same offspring distribution had been sampled more intensively.

### Age for defining a cohort

Consider all *N*_*1*_ members of a cohort that reach age 1, and their production of offspring that also survive to age 1. At stable *N*,

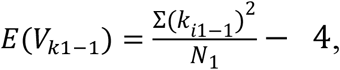

and making this substitution in Equation 1 produces

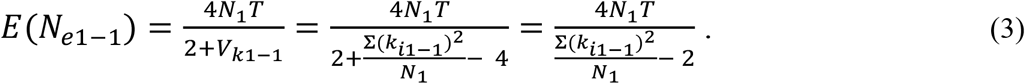

We want to compare this with *E*(*N*_*eα*−*α*_) for the cohort defined at age α. The adult cohort size is reduced by cumulative mortality through age at maturity (*l*_*α*_), so the cohort size of individuals that reach reproductive age is *N*_*α*_ = *l*_*α*_*N*_*1*_, and

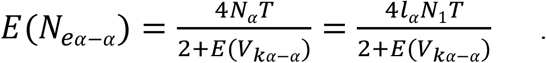

We use a two-step process to find *E*(*V*_*kα−α*_) as a function of parameters for newborns, for comparison with Eq. 3. First, consider production of newborn offspring by adults that survive to age α (*k*_*iα*−1_). The only members of the newborn cohort with *LRO*>0 are those that survive to adulthood, and these are the same individuals that make up the adult cohort. Therefore, the following identities hold:

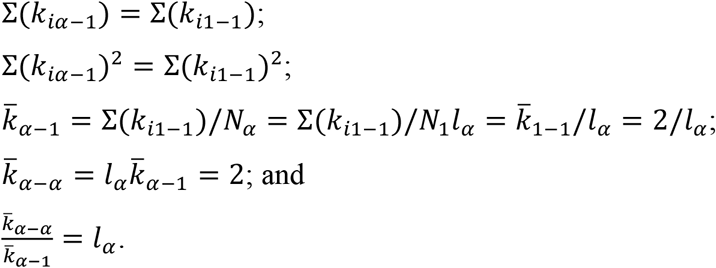

Now we have an expression for the variance of newborn offspring produced by members of the age α cohort:

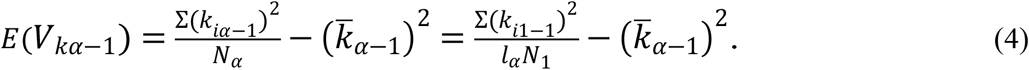

Because 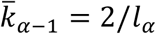 is larger than required to produce a stable population, we do Crow-Morton rescaling to obtain *E*(*V*_*kα−α*_) at stable *N*. A key component of the rescaling is the raw variance-to-mean ratio, which is

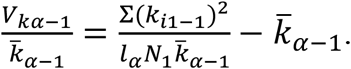

Substituting for 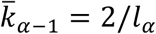:

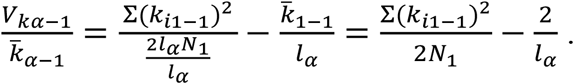

Now we use Equation 2b to rescale the raw variance *V*_*k*α−1_ to its expected value at 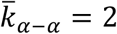:

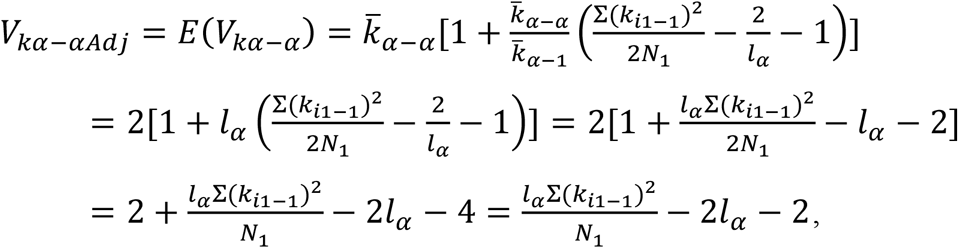

which leads to

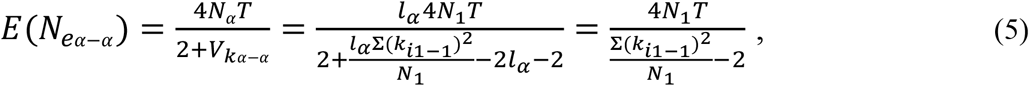

which is identical to Equation 3.

### Non-random survival

The above analyses assumed random survival of newborns to age at maturity. But what if survival is not random, even though mean survival rate remains the same? Survival can be non-random in a variety of ways, but for variance in offspring number and hence *N*_*e*_, the only thing that matters is whether survival is family-correlated, such that offspring from different families have different probabilities of survival. Crow and Morton (1955) provided an analytical solution for one extreme form of family-correlated survival, where entire families either survive or not as a unit. Under those conditions, based on Equation 4 in Waples (2002) and using the notation in Eq 2b:

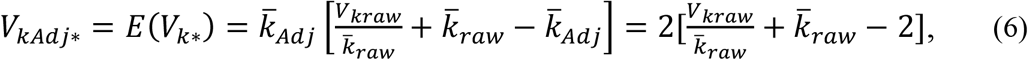

where * indicates a result expected under family-correlated survival.

Consider first a cohort defined as newborns. There are *N*_*1*_ individuals, each representing one potential parent. However, all offspring are produced by the *N*_*α*_ individuals that survive to maturity, and the distribution of offspring number is not affected by which individuals survive to adulthood. Therefore, if the cohort is defined in terms of newborns, *N*_*e*1−1_ calculated using Equation 3 is the same regardless which individuals survive to adulthood.

The situation is different when *N*_*e*_ is calculated for adults producing adults. In this case, the cohort has *N*_*α*_ = *l*_*α*_*N*_*1*_ individuals, and *E*(*V*_*k*α−1_) is given by Equation 4. Using Eq 6 to predict *V*_*kα−α*_ under family-correlated survival:

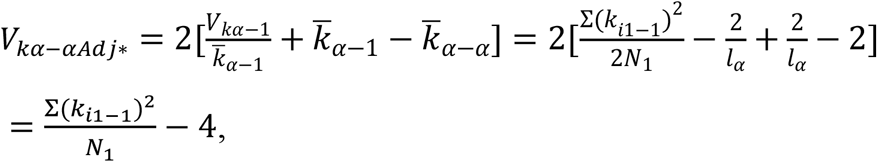

and

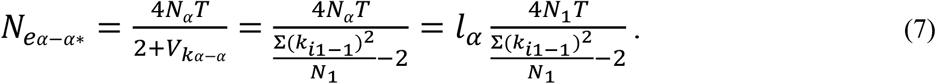

In this extreme form of family-correlated survival, provided that reproduction is measured in terms of adults producing adults, *N*_*e*_ is reduced by the factor *l*_*α*_ compared to the result for random survival to adulthood.

### Null parents

Finally, we want to consider how null parents (those with *LRO* = 0) affect Hill’s *N*_*e*_. For simplicity, we focus on newborn cohorts, and for comparative purposes we use results for the full cohort (including nulls) of *N*_*1*_ individuals, for which (from Equation 3),

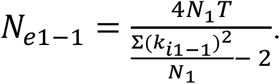

Null members of the newborn cohort can achieve *LRO* = 0 in two different ways: (1) with probability *l*_*α*_, they can fail to survive to age at maturity, or (2) they can reach age at maturity but never produce any offspring. Here we consider a scenario where *LRO* data are available for all individuals with *LRO*>0 (and perhaps for some null parents), but not for at least a subset of null individuals. The subscript ‘*x*’ will be used for variables associated with the *N*_*x*_ individuals for which *LRO* data are available. Then λ = *N*_*x*_/*N*_*1*_ is the fraction of the newborn cohort for which data are available, and the number of null parents without *LRO* data is *N*_*null*_ = *N*_*1*_–*N*_*x*_ = *N*_*1*_(1-λ). The objective is to find *N*_*ex*_ and compare that with *N*_*e1-1*_ as calculated above.

Null parents do not affect Σ(*k*_*i*1−1_) or Σ(*k*_*i*1−1_)^2^, so mean and variance of *LRO* (in terms of production of newborns) for the *N*_*x*_ individuals are

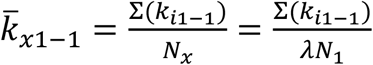

and

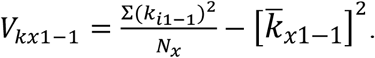

Because 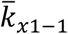 is >2, we do Crow-Morton rescaling assuming random survival to find *V*_*kx*1−1*Adj*_ when 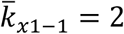. From Equation 2b,

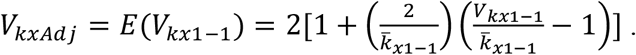

Substituting for 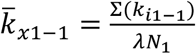:

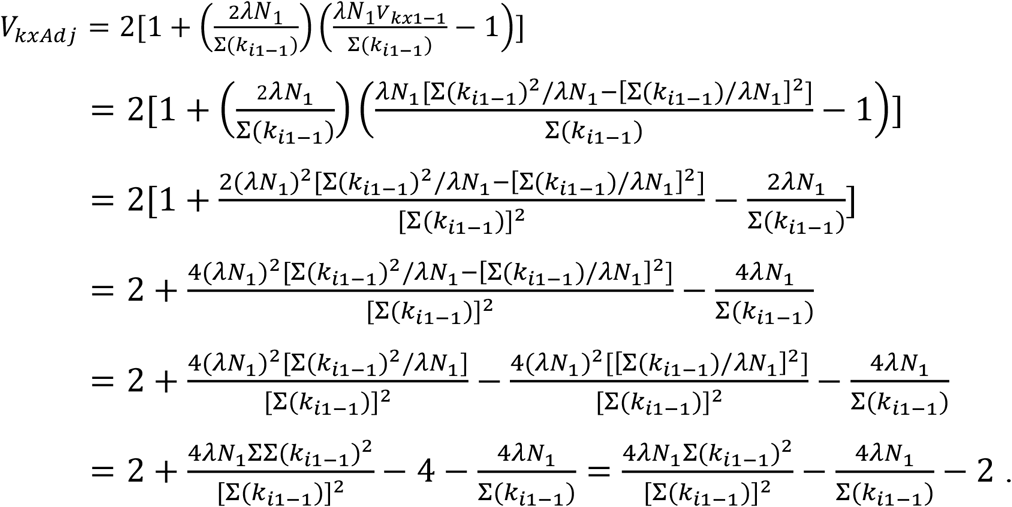

Therefore,

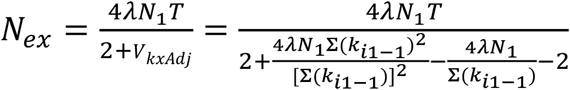

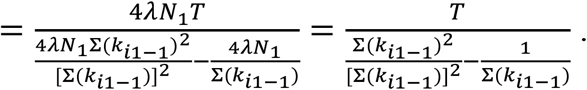

Substituting for Σ(*k*_*i*1−1_) = 2*N*_1_:

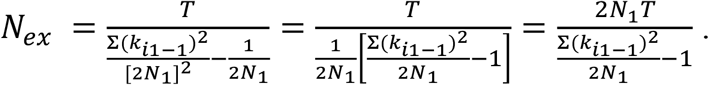

Multiplying the top and bottom by 2 produces

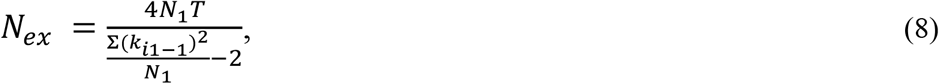

which is identical to the result (Equation 3) found using all *N*_*1*_ members of the cohort, including nulls.

### Computer simulations

The analytical results obtained above were checked using computer simulations. All coding was done in R v 2.2.2 (R Core Team 2022). The model species had 10 age classes and constant survival probability of *s*_*x*_ = 0.7/year. ‘Newborns’ were age 1, and both sexes matured at age α = 3. Females had constant fecundity with age and, within ages, Poisson variance in reproductive success. In males, fecundity was proportional to age, and within ages variance in offspring number was overdispersed (being on average twice the mean). In both sexes, age-specific fecundity (*b*_*x*_) was scaled to values expected to produce a stable population (Table 1).

**Table 1.** Age-specific vital rates for the model species used in the simulations. Age at maturity is α = 3 and maximum age is 10. Survival is constant in both sexes at *s*_*x*_ = 0.7/year, and the vector *l*_*x*_ is cumulative survival through age *x*. Age-specific fecundity (*b*_*x*_) is the expected number of offspring produced in one year by an individual of age *x*; it is scaled to values that will produce a stable population. *ϕ*_*x*_ is the ratio of variance to mean offspring number within ages. In females, *b*_*x*_ is constant with age, and within each age the distribution of offspring number is Poisson (all *ϕ*=1). In males, *b*_*x*_ is proportional to age, and within each age variance in offspring number is twice *b*_*x*_, so all *ϕ*=2.

| Age | Females |  |  |  | Males |  |  |  |
| --- | --- | --- | --- | --- | --- | --- | --- | --- |
| | $s_x$ | $l_x$ | $b_x$ | $\phi_x$ | $s_x$ | $l_x$ | $b_x$ | $\phi_x$ |
| 1 | 0.7 | 1 | 0 | 0 | 0.7 | 1 | 0 | 0 |
| 2 | 0.7 | 0.70 | 0 | 0 | 0.7 | 0.70 | 0 | 0 |
| 3 | 0.7 | 0.49 | 1.30 | 1 | 0.7 | 0.49 | 0.80 | 2 |
| 4 | 0.7 | 0.34 | 1.30 | 1 | 0.7 | 0.34 | 1.07 | 2 |
| 5 | 0.7 | 0.24 | 1.30 | 1 | 0.7 | 0.24 | 1.34 | 2 |
| 6 | 0.7 | 0.17 | 1.30 | 1 | 0.7 | 0.17 | 1.61 | 2 |
| 7 | 0.7 | 0.12 | 1.30 | 1 | 0.7 | 0.12 | 1.88 | 2 |
| 8 | 0.7 | 0.08 | 1.30 | 1 | 0.7 | 0.08 | 2.15 | 2 |
| 9 | 0.7 | 0.06 | 1.30 | 1 | 0.7 | 0.06 | 2.41 | 2 |
| 10 | 0 | 0.04 | 1.30 | 1 | 0 | 0.04 | 2.68 | 2 |

Newborn sex ratio was 1:1, and each year the adult population produced a total of *N*_1_ = 100 yearling offspring of each sex. Subsequently, stochastic variation in survival rates allowed the realized sex ratio and age structure to vary. Reproduction followed the generalized Wright-Fisher model described by Waples (2020, 2022), in which parents are allowed to have unequal probabilities of producing offspring, with these probabilities being considered ‘parental weights.’ Fecundity was fixed in females with Poisson variance, so each year all adult females behaved like a Wright-Fisher population with equal expectations of reproductive success (hence all parental weights were the same). Because by chance some females lived longer and had more years to accumulate *LRO*, variance in lifetime offspring number in females still exceeded the mean (which on average was 2). In males, older individuals were more likely to be chosen as the parent of a given offspring. Furthermore, because male within-age variance was overdispersed, an additional random component was added to the parental weights. This random component was generated independently every year, so there were no persistent individual differences in reproductive success; that is, except by chance, no individuals were consistently above or below average for their age and sex at producing offspring. Furthermore, reproduction and subsequent survival were uncorrelated over time. Under those conditions, given the age-specific vital rates shown in Table 1, the software AgeNe (Waples et al. 2011) can be used to predict generation length, *V*_*k*_, and *N*_*e*_ separately for each sex.

After initialization to establish the expected age structure determined by the *l*_*x*_ vector, each simulation was run for 500 years. Each year, newborn cohorts of *N*_1_ males and *N*_1_ females were generated, and each cohort provided a separate opportunity to estimate *N*_*e*_, which was done separately for males and females. A Pedigree file kept track of the male and female parents and their ages for each offspring. *LRO* for each member of a newborn cohort was calculated by counting the number of subsequent offspring having that individual as a parent. One lifespan (10 years) of data were dropped at the start and end of the time series, and results were averaged over the remaining 480 years. The core analyses modeled random survival of newborns to adulthood (age 3) by randomly choosing survivors with probability *l*_*α*_ = *l*_3_ = 0.7^2^ = 0.49. A separate file kept track of ID numbers of survivors to age 3, and this information was used with the cohorts defined in terms of age 3 adults to ensure that their offspring were only recorded if they also survived to age 3. Non-random survival of juvenile offspring was modeled by selecting entir families to either survive to adulthood or not.

## RESULTS

Hill’s model defines cohorts in terms of newborns. Assuming constant *N*, random survival of newborns, and independence of survival and reproduction as modeled here, the software AgeNe predicts generation length, *V*_*k*_, and *N*_*e*_ for both sexes based on the age-specific vital rates in Table 1. As shown in the top section of Table 2, in both sexes, mean generation length and *V*_*k*_ and harmonic mean 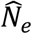 were nearly identical to the AgeNe expectations. Expected *N*_*e*_ is lower for males than females (125 vs 159) because increasing fecundity with age and overdispersed within-age variance combine to substantially increase lifetime variance in reproductive success (E(*V*_*k*_) = 15.9 for males vs 10.2 for females).

**Table 2.** Simulation results for both sexes, for cohorts defined as newborns and adults. Expected results (Exp) are from AgeNe (Waples et al. 2011); observed results (Obs) are means across 480 replicate cohorts. *N* = cohort size; *N*_*null*_ = number of cohort members with *LRO* = 0; *k*_*max*_ = maximum *LRO* within a cohort;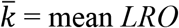; *V*_*k*_ = raw var(*LRO*); *V*_*kAdj*_ = rescaled *V*_*k*_; *T*= generation length;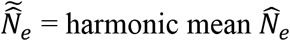. The ‘null’ subscript indicates results that excluded all null parents.

| | | | $N$ | $N_{null}$ | $k_{max}$ | $\bar{k}$ | $V_k$ | $V_{kAdj}$ | $T$ | $\tilde{N}_e$ | $\bar{k}_{null}$ | $V_{knull}$ | $V_{knullAdj}$ | $\tilde{N}_{enull}$ |
| --- | --- | --- | --- | --- | --- | --- | --- | --- | --- | --- | --- | --- | --- | --- |
| Newborns |  |  |  |  |  |  |  |  |  |  |  |  |  |  |
| Male | Exp |  | 100 | - | - | 2.00 | 15.9 | 15.9 | 5.59 | 125 | - | - | - | 125 |
|  | Obs |  | 100 | 64.2 | 20.4 | 2.00 | 15.7 | 15.9 | 5.59 | 125 | 5.60 | 23.8 | 4.30 | 125 |
| Female | Exp |  | 100 | - | - | 2.00 | 10.2 | 10.2 | 4.84 | 159 | - | - | - | 159 |
|  | Obs |  | 100 | 56.0 | 14.2 | 2.00 | 10.1 | 10.2 | 4.83 | 158 | 4.55 | 11.3 | 3.29 | 158 |
| Adults |  |  |  |  |  |  |  |  |  |  |  |  |  |  |
| Male | Exp |  | 49 | - | - | 2.00 | - | - | 5.59 | 125 | - | - | - | 125 |
|  | Obs |  | 49.3 | 19.1 | 10.9 | 2.01 | 6.59 | 6.72 | 5.59 | 125 | 3.26 | 6.72 | 3.25 | 126 |
| Female | Exp |  | 49 | - | - | 2.00 | - | - | 4.84 | 159 | - | - | - | 159 |
|  | Obs |  | 49.0 | 12.3 | 8.1 | 2.03 | 3.82 | 3.91 | 4.83 | 158 | 2.68 | 3.43 | 2.36 | 160 |

The remainder of the results are discussed below with respect to the four issues outlined in the Introduction.

### Mean offspring number ≠ 2

In a population of constant size, mean offspring number must be 2. Using data with 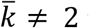 to compute *N*_*e*_ using Equation 1 can affect both bias and precision, depending whether the departures from 2 are random or systematic. Systematic bias is most easily demonstrated by considering reproductive success data for only part of the life cycle—for example, production of yearling offspring by adults. This scenario was modeled analytically in Methods, with subscripts ‘α-1’, indicating production of age-1 offspring by cohorts of individuals that reached age α (hence adults). With *N*_1_ = 100 newborns of each sex, the fraction *l*_3_ = 0.49 = 49 individuals are expected to survive to adulthood, and their expected mean *LRO* is 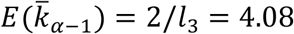, which is more than twice the value needed for a stable population. Because of the positive correlation between *V*_*k*_ and 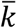, raw *V*_*k*_ for juvenile offspring produced by adults will be inflated, which will cause a downward bias in estimating variance *N*_*e*_. This is the exact scenario the Crow-Morton rescaling method was designed to deal with. Rescaling the inflated raw *V*_*k*_ shrinks it to the value expected if 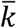 had been 2.0, which would then model production of adults by adults. In the simulations using adult male cohorts (adults producing adults; bottom half of Table 2), mean 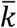 and *V*_*k*_ and harmonic mean 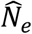 were 2.00, 6.72, and 125, respectively. Using the raw simulated data for male adults producing yearling offspring, the comparable values were mean 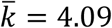, mean *V*_*k*_ = 23.3, and harmonic mean 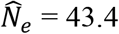 – the latter being just over a third of the true value. After rescaling, mean *V*_*kAdj*_ (6.81) and harmonic mean 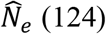 were close to the expected values for a full lifecycle. Results were qualitatively similar for females: after rescaling, mean raw *V*_*k*_ dropped from 11.6 to 3.97 (close to the 3.91 value found in simulations of adults producing adults), and harmonic mean 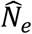 increased from 69.5 to 156.7 (close to the expected value of 159). The pronounced difference between 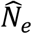 calculated on scaled vs raw data is apparent for both sexes in Figure 2.

**Figure 1.**
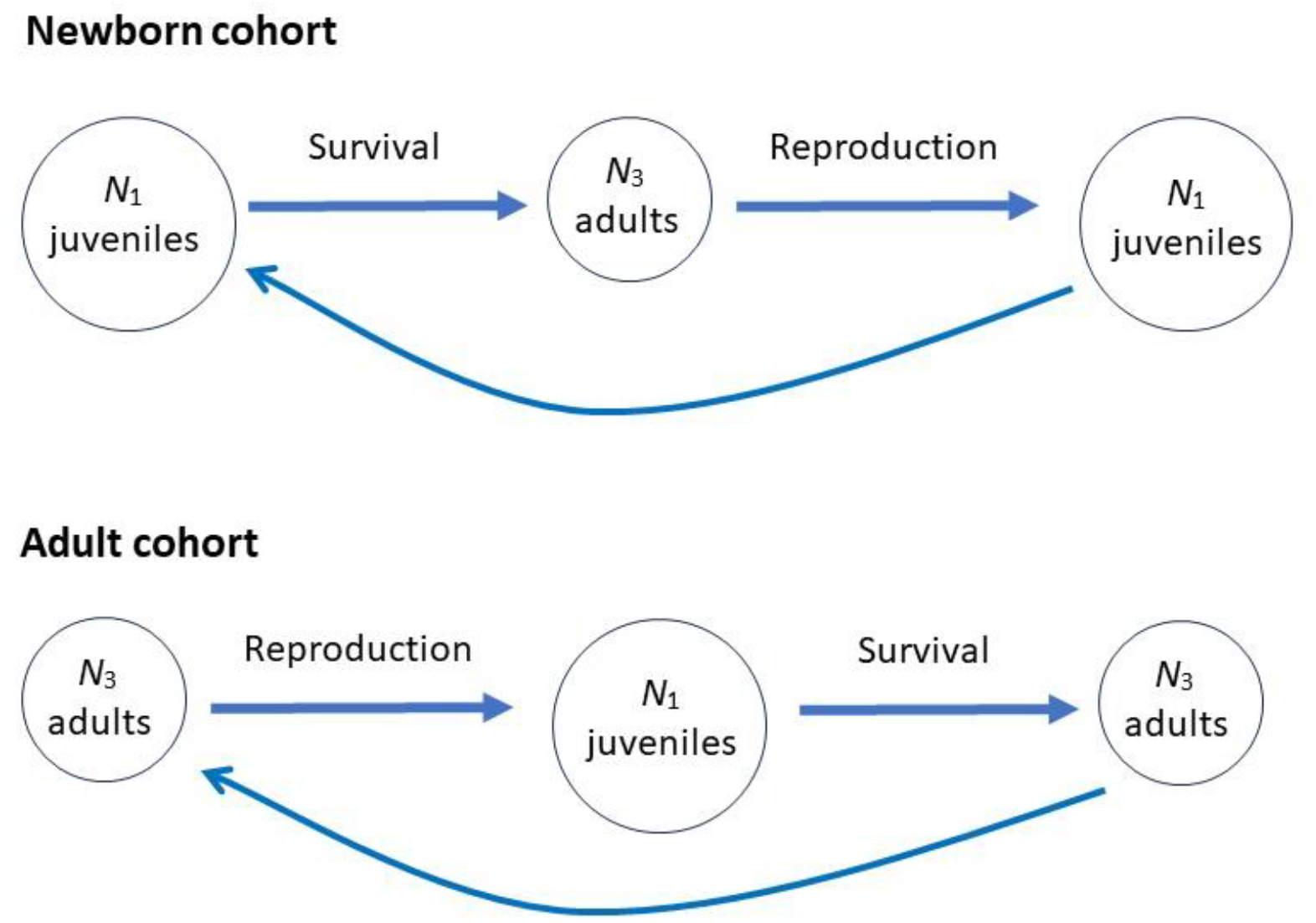
Schematic diagram illustrating the difference between defining cohorts in terms of newborns (age 1) and adults (age 3). *N* is the number of individuals alive at the indicated age.

**Figure 2.**
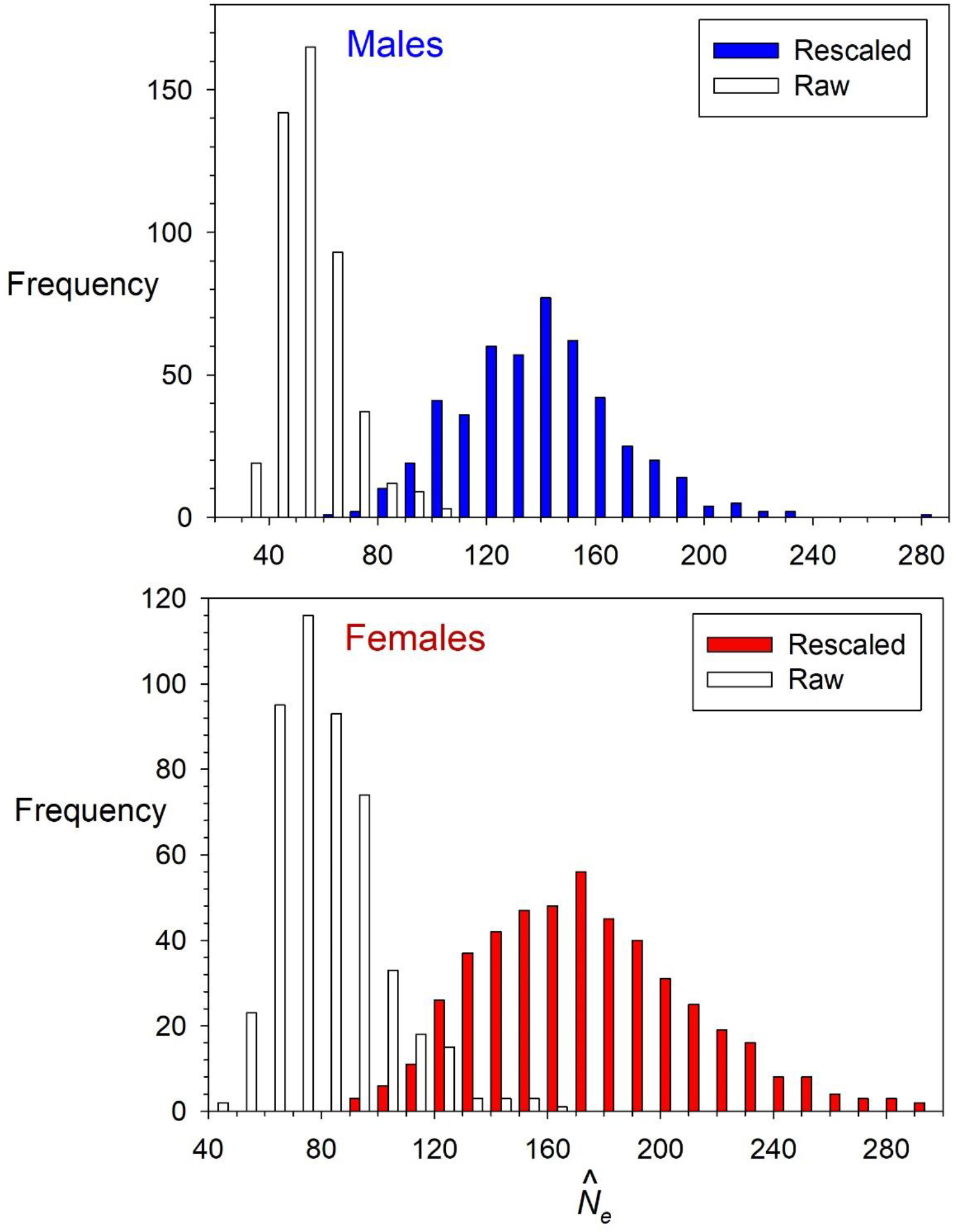
Distribution of 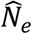 for single cohorts in simulated data. In this scenario, the cohort is defined in terms of age-3 adults but offspring are counted at age 1, so only part of a lifecycle is covered by the data. If raw *V*_*k*_ is used to estimate effective size, 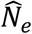 is biased downwards (open bars); when raw *V*_*k*_ is rescaled using Equation 2b (filled bars), the harmonic mean 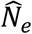 (123.8 for males and 156.7 for females) agreed closely with results for adults producing adults (Table 2, bottom, harmonic mean 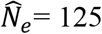 for males and 158 for females).

In the simulations using newborn cohorts, the empirical 95% confidence interval for 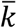 was 1.38-2.64 for males and 1.37-2.76 for females, with both means being essentially identical to the expected 2.0. This random variation around the stable-*N* expectation also affects *V*_*k*_ and hence 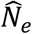 in each cohort, but it does not lead to appreciable bias because the deviations in mean offspring number are approximately symmetric around 2.0. However, rescaling *V*_*k*_ in each replicate does substantially increase precision (Figure 3). After rescaling, the coefficient of variation (CV) for male 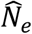 dropped from 0.25 to 0.16, and for females the CV dropped from 0.19 to 0.14 (Figure 3).

**Figure 3.**
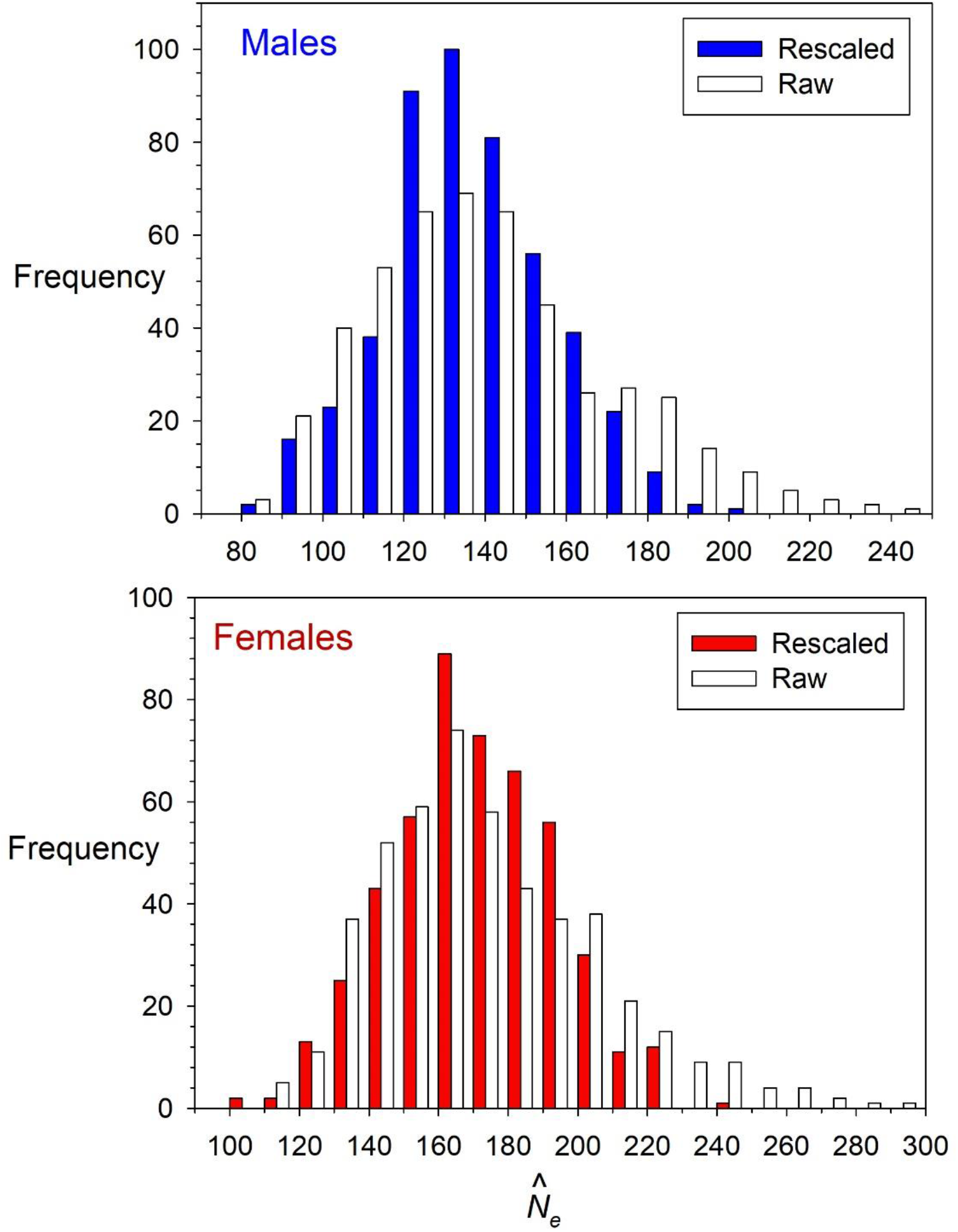
Distribution of 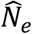 for newborn cohorts in simulated data. Open bars show results when raw *V*_*k*_ is used to estimate effective size; filled bars show results when raw *V*_*k*_ is rescaled to its expected value at 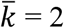 using Equation 2b. Harmonic mean 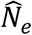 across replicates was not affected by rescaling (in both cases being 125 for males and 158 for females; Table 2), but rescaling substantially reduces 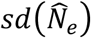.

Rescaling does not eliminate all random demographic stochasticity inherent to finite populations. However, given the underlying assumption of stable *N* and stable Age structure, different cohorts provide replicate estimates of the same parametric *N*_*e*_, which means that precision can be increased by averaging results from multiple cohorts. Statistical theory indicates that the standard deviation of the mean of a random variable is inversely proportional to the square root of the number of values across which the mean is calculated. With respect to *N*_*e*_ estimation, this means that

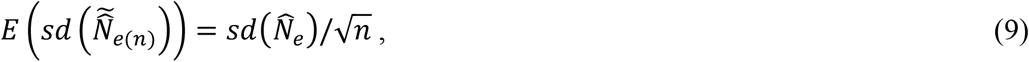

where 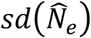 is the standard deviation of estimates based on a single cohort and 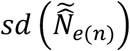 is the standard deviation of the composite estimates, each of which is the harmonic mean of *n* single-cohort estimates. When random resampling was used to generate many replicate estimates of 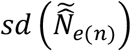, the increase in precision for larger values of *n* closely agreed with predictions from Equation 9 (Figure 4).

**Figure 4.**
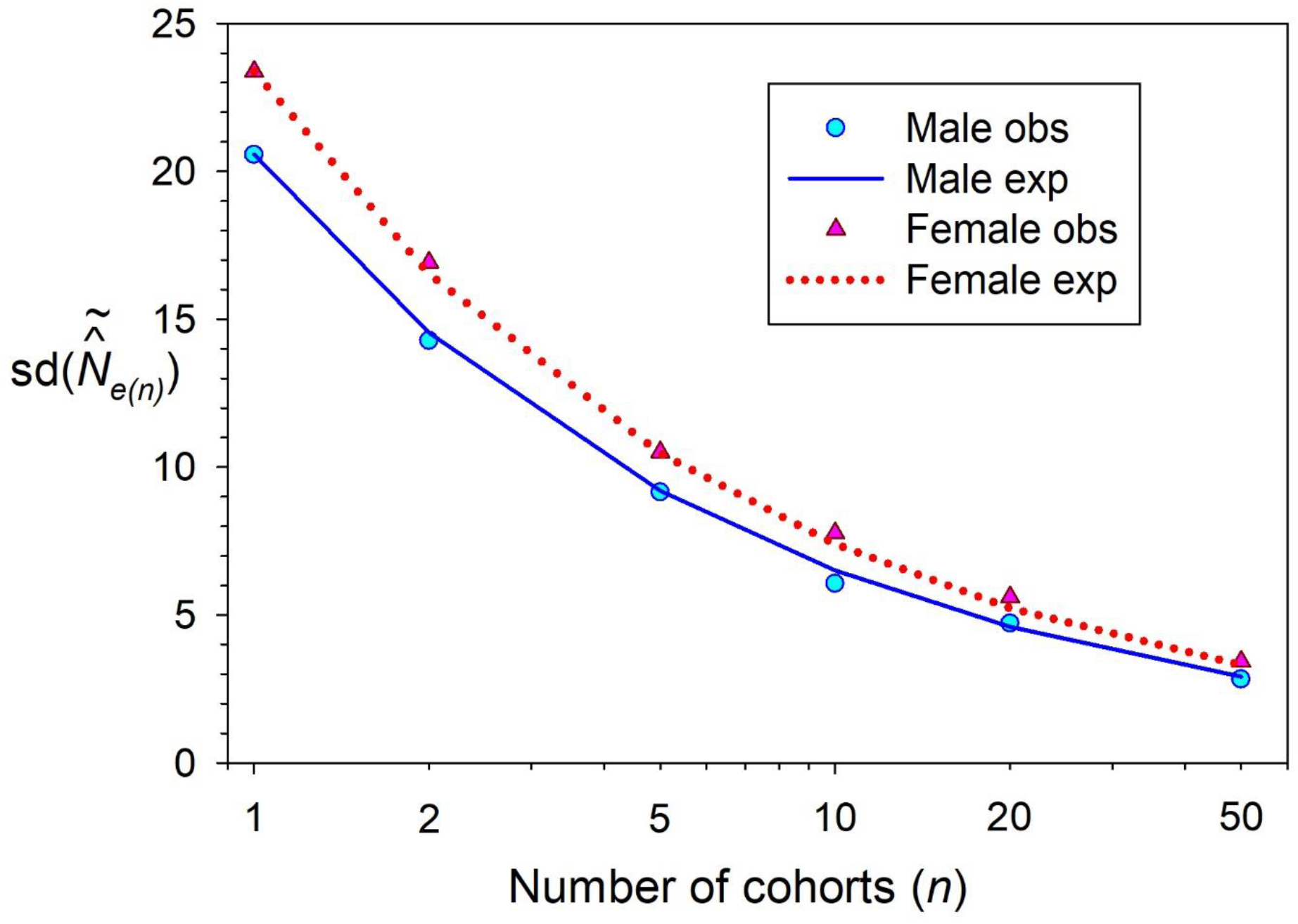
Reduction in the standard deviation of harmonic mean 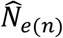 as data from more replicate cohorts are used in the computation. Symbols (obs) are results from the simulations for newborn cohorts; lines (exp) are expectations based on Equation 9. Note the log scale on the *X* axis.

### Age for defining a cohort

Analytical results in Methods showed that, if survival of newborns to adulthood is random, 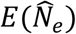 is the same for cohorts defined in terms of *N*_1_ newborns or *N*_α_ adults (compare Equations 3 and 5). The cohort size (which appears in the numerator of Equation 1) is smaller when it is defined in terms of adults, but this is exactly offset by a reduction in *V*_*k*_, which appears in the denominator. The simulations confirmed these analytical predictions (Table 2). Mean *V*_*kAdj*_ was smaller for adult cohorts (6.72 vs 15.9 for newborn cohorts in males; 3.91 vs 10.2 in females), but harmonic mean 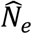 was essentially the same (125 in males and 158 in females, regardless how the cohort was defined).

Age at which a cohort defined can, however, affect precision. Cohorts defined as adults are smaller (for the model species used in the simulations, an average of 49% the size of a newborn cohort), which leads to more random sampling error in estimating parameters. In males, 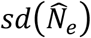 was 29.6 for adult cohorts compared to 20.6 for newborns (an increase of 44%), and in females the increase was proportionally larger (61%, 37.7 for adults vs 23.4 for juveniles).

### Non-random survival of juveniles

Although cohort age does not affect bias when juvenile mortality is random, that is not the case when different families have different probabilities of survival from newborns to adulthood. The consequences of family-correlated survival are missed when cohorts are defined in terms of newborns, where mortality occurs before reproduction (Figure 1). With random survival, each newborn either survives to adulthood or not, with probability *l*_*α*_. With family-correlated survival, the result is exactly the same: each newborn represents a separate potential family of size 1, which then survives or not to adulthood, with probability *l*_*α*_. The situation is different for cohorts defined in terms of adults, in which case reproduction occurs first and a range of different family sizes is generated prior to the episode of family-correlated survival.

When each family either survives or not as a unit, *E*(*N*_*e*_) is reduced by the factor *l*_*α*_ (Equation 7), which is just the juvenile-adult survival rate. In a simulation of this non-random survival scenario, harmonic mean 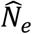 was 64.8 for males and 80.7 for females, both close to the expectations of 61.6 and 77.9 based on Equation 9. These results apply to an extreme form of family-correlated survival, so the magnitude of the effect would be smaller for weaker forms of viability selection. Hill (1972) stated without proof that, although his model assumed cohorts defined in terms of newborns, bias associated with viability selection of offspring could be avoided by defining cohorts in terms of adults, and results presented here support that statement.

### Null parents

If some or all members of a cohort that are null parents (having *LRO* = 0) are not included in the empirical data, 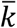, *V*_*k*_ and the nominal cohort size all are affected, but these variables change in such a way that 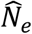 does not change. This result was predicted analytically in Methods (compare Equations 3 and 8) and confirmed in the simulations (Table 2): for both newborn and adult cohorts, excluding nulls increased 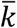 and raw *V*_*k*_ but had no effect on 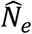. This parallels the results for a discrete-generation model reported by Waples and Waples (2011), who showed that the conclusion that null parents do not affect *N*_*e*_ holds generally for inbreeding *N*_*e*_, but for variance *N*_*e*_ it is true only for stable *N* and 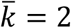. Under Hill’s stable-*N* model, inbreeding and variance *N*_*e*_ must be the same, which explains why the null-parent result holds for his overlapping-generation model.

## Discussion

A key parameter in Hill’s method (Equation 1) is *V*_*k*_ = the variance in lifetime number of offspring produced by members of a single cohort (that is, individuals born in the same year or season). Recent technological advances in (a) speed and affordability of DNA sequencing and (b) ability to obtain high-quality DNA non-lethally make it increasingly feasible to reconstruct pedigrees in the wild (Pemberton 2008; Riester et al. 2009; Huisman 2017). Nevertheless, tracking reproductive output across individual lifetimes remains very challenging, especially for long-lived species. As a consequence, achievable experimental designs and empirical parameter estimates from fieldwork rarely conform exactly to assumptions of models. Results related to the four potentially problematic scenarios described in the Introduction can be summarized as follows.

### Mean offspring number ≠ 2

When mean offspring number in a sample is not 2, rescaling *V*_*kraw*_ to its expected value at 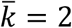 using Equation 2B will improve performance of Hill’s method. The nature of the improvement depends on whether departures from 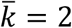 are systematic or random. If the population as a whole is dynamically stable, measurements of *LRO* are comprehensive and cover a full lifecycle, and departures from 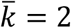 reflect only demographic stochasticity and random sampling error, then using *V*_*kraw*_ to estimate *N*_*e*_ in Equation 1 will not lead to bias, but the variance of 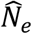 will be higher than it would if each *V*_*kraw*_ were rescaled (Figure 3).

Two general scenarios lead to systematic departures from 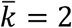, and these lead to bias in 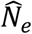 if *V*_*kraw*_ is used without rescaling. First, if the measurement of *LRO* does not cover a full life cycle (for example, if the cohort is defined as adults but offspring are enumerated as juveniles), it generally will be the case that 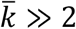. In that case, *V*_*kraw*_ will be systematically biased upwards and the resulting 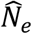 will be biased downwards, as in Figure 2. Second, even if the population is stable and estimated *LRO* spans a full lifecycle, incomplete sampling of offspring (or inability to match all offspring to parents) will cause 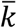 to be consistently less than 2. This will lead to an underestimate of true *V*_*k*_ and an upward bias in 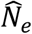. Rescaling *V*_*kraw*_ using Equation 2a eliminates the bias for both scenarios.

### Age for defining a cohort and juvenile survival

These two factors are considered together here because they interact. If survival from juvenile to adult stages is random, then it does not matter at what age the cohorts are defined, provided that measurement of *LRO* spans a full life cycle. The same is not true for non-random survival: in that case, 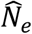 will be biased upwards if newborn cohorts are used, because in that case the consequences of non-random survival are missed. As noted by Hill (1972, p 285). this bias can be avoided by defining cohorts in terms of adults: “A familial correlation of viability to adulthood can be taken into account by specifying the number of adult individuals entering the population each generation, and the variance of the number of adult progeny per adult individual.” However, Hill only mentioned this key point in one sentence of his 1972 paper, and not at all in his 1979 version, so its significance has likely been underappreciated.

It is important to clarify what type of non-randomness in survival is important with respect to Hill’s method. Individuals can be characterized according to a variety of phenotypic traits (e.g., size, coloration, physiology, behavior, etc.), and survival potentially could be correlated with any of these traits without affecting 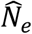. The only thing that matters is whether the probability an individual survives or not depends on which family it comes from (and hence who its parents are). If so, survival is family correlated, which means that siblings do not have independent probabilities of surviving. Family-correlated survival can arise from heritable traits passed on from parents to offspring, but it also commonly happens for non-genetic reasons early in life history, when family members co-occur in space and time and can be jointly affected by random events, such as storms, drought, or nest predation. If cohorts are defined in terms of adults, any family-correlated survival that does occur is reflected in the variance of *LRO* among adults, but that is not the case for newborn cohorts.

As noted above, if *LRO* is measured for only part of a life cycle (adults producing adults), the expected result is that 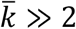 and *V*_*kraw*_ will be upwardly biased. Three general scenarios can be considered, depending on the objectives and the species’ biology.

1. The objective is to estimate *N*_*e*_ for the part of the life cycle for which data are available. In that case, rescaling to 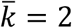 using the Crow-Morton method is appropriate, since it removes the bias associated with large mean offspring number. Nothing is inferred regarding survival to adulthood, as the estimate only applies to the adult-juvenile phase.
2. The objective is to estimate *N*_*e*_ for the full life cycle, and random survival until adulthood can be assumed. In that case, calculation of 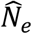 is the same as in scenario (1), the only difference being that 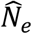 is interpreted as applying to a full generation rather than a fraction of a generation.
3. The objective is to estimate *N*_*e*_ for the full life cycle, and random survival until adulthood cannot be assumed. This is the most challenging scenario, as 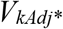 and 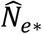 depend on accurately characterizing the magnitude of the effects of family-correlated survival. Rescaling *V*_*kraw*_ using Equation 6 implements an extreme form of family-correlated survival that would likely overestimate 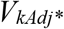 and underestimate 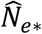 in most realistic scenarios. Additional research and modeling is needed to more fully explore the range of plausible survival scenarios.

### Null parents

Null parents occur naturally in most populations, as some individuals by chance produce no offspring that survive to age at enumeration. But these individuals also can have characteristics that affect their probability of being sampled; for example, if they don’t participate in reproduction, they can be invisible to sampling efforts and likely will not be recorded as potential parents. Therefore, it is important to consider how omitting null parents from calculations of cohort size, 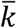, and *V*_*k*_ might affect bias. The good news is that omitting some or all individuals with *LRO* = 0 has no effect on 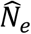, provided that the raw data are scaled to expectations for 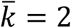.

### Sampling multiple cohorts

Even with complete *LRO* data spanning a full life cycle in a stable population, the estimate of *N*_*e*_ from a single cohort is inherently stochastic (Figure 3). However, if the *N*_*e*_/*N* ratio is also constant, samples from multiple cohorts provide independent estimates of the same parameters, which can considerably reduce uncertainty of the composite estimates Figure 4).

## Acknowledgments

I am grateful to Joachim Mergeay for useful discussions. The author declares no competing interest.

## Data Archiving

All results presented here were generated by computer simulations. R code to conduct the simulations will be provided on acceptance.

